# Routing States Transition During Oscillatory Bursts and Attentional Selection

**DOI:** 10.1101/2022.10.29.514374

**Authors:** Kianoush Banaie Boroujeni, Thilo Womelsdorf

## Abstract

Neural information routing relies on spatiotemporal activity dynamics across interconnected brain areas. However, it remains unclear how routing states emerge at fast spiking timescales and interact with the slower activity dynamics of larger networks during cognitive processes.

Here, we show that localized neural spiking events generate long-range directional routing states with spiking activity in distant brain areas that dynamically switch or amplify during oscillatory bursts, selective attention, and decision-making. Computational modeling and neural recordings from lateral prefrontal cortex (LPFC), anterior cingulate cortex (ACC), and striatum of nonhuman primates revealed that cross-areal, directional routing states arise within ∼20 ms around spikes of single neurons. On average, LPFC spikes led activity in the ACC and striatum by few milliseconds. The routing state was amplified during LPFC beta bursts between the LPFC and striatum and switched direction during ACC theta/alpha bursts between ACC and LPFC. Selective attention amplified the lead of these theta/alpha-specific lead-ensembles in the ACC, while decision-making amplified the lead of ACC and LPFC spiking output over the striatum. Notably, the fast lead/lag relationships of cross-areal neuronal ensembles that were modulated by attention states or decision-making predicted firing rate dynamics of their neurons during those functional states at slower timescales. Overall, our findings demonstrate directional routing of spiking activity across nonhuman primate frontal and striatal areas, as well as the functional and network states that modulate the direction and magnitude of these interactions.

**Summary:** Fast spatio-temporal dynamics of brain activity subserves the routing of information across distant regions and is integral to flexible cognition, decision-making, and selective attention. This study demonstrates that routing dynamics emerge as 20 ms brief lead and lag relationships of spiking activities across distant brain areas. The direction and magnitude of the lead and lag relationships systematically switched during frequency-specific oscillatory bursts and when attention shifts to visual cues.

## Introduction

Routing states during flexible behaviors depend on precise spatio-temporal dynamics of neuronal activity across interconnected brain areas (Singer, 2013). These dynamics proceed at varying temporal scales ranging from rapid synaptic communication (Markram et al., 1997; Hasenstaub et al., 2005), fast recurrent inhibitory and excitatory neuronal interactions (Hahn et al., 2014 p.20; Palmigiano et al., 2017), and slower firing rate modulation in neural populations (Murray et al., 2014; Luczak et al., 2015). As for spatial scales, they can range from a single neuron to multiple nearby neurons, a local population of neurons, or a distributed network of neurons across distant brain areas (Einevoll et al., 2013; Lewis et al., 2015; Wang, 2022). However, the relationship between these temporal and spatial scales during dynamic transitions of brain-wide routing states has remained unclear.

To understand the spatio-temporal dynamics of brain-wide routing states during flexible behaviors, we need to evaluate the directionality of sequential spiking activity in neuronal ensembles distributed in distant brain areas. Achieving this goal remains challenging which is evident by a lack of direct insights about long-range, directional spiking interactions at fast timescales in behaving animals. First, fast activity coordination of neural activity in extracellular recordings has been measured directly by cross-correlating spike trains or indirectly quantifying spike synchronization to LFP in the gamma frequency band (Womelsdorf et al., 2007; Smith and Kohn, 2008), but these methods are often not reliable and robust in estimating the direction of spiking interactions. Second, attempts to measure directionality in spiking networks, thus far, have largely focused on latent variables in higher-order correlational structures of spiking dynamics over pre-defined neural populations without providing mechanistic insights about fast directional interactions (Semedo et al., 2019; van Kempen et al., 2021). Third, studying long-range directional communication between cortical and subcortical areas subserving higher cognitive functions in the primate and nonhuman primate brain has been limited to spike-LFP and LFP-LFP measures which provide only indirect insights about cross-areal spike-to-spike directional interactions within neuronal ensembles (Gregoriou et al., 2009; Salazar et al., 2012; Siegel et al., 2012; Antzoulatos and Miller, 2014; Womelsdorf et al., 2014; Fries, 2015; Fiebelkorn et al., 2018).

Here, we overcome these limitations and unravel the spatio-temporal dynamics of neural activities within and across brain areas of the medial and lateral fronto-striatal network. Connectivity of prefrontal and striatal areas is hierarchically organized and rapidly engages and disengages during sequential and flexible behaviors (Jin et al., 2009; Haber and Knutson, 2010). Key nodes of this network are the anterior striatum (head of the caudate nucleus), lateral prefrontal cortex (LPFC, area 49/9), and anterior cingulate cortex (ACC, area 24) (Ferry et al., 2000; Haber and Knutson, 2010). Activity changes in each of these areas correlates with learning success as evident at the levels of single neuron firing (Antzoulatos and Miller, 2011; Oemisch et al., 2019; Boroujeni et al., 2020; Banaie Boroujeni et al., 2021) as well as oscillatory bursting of the local field potential (LFP) (Feingold et al., 2015; Lundqvist et al., 2016). In addition, activity of single neurons in the LPFC, ACC, and striatum synchronizes to narrow-band frequency fluctuations of the LFP during flexible behavior and efficient attentional control (Antzoulatos and Miller, 2014; Womelsdorf et al., 2014; Voloh et al., 2020). These findings have established that single neuron spiking and network-level LFP activity in the fronto-striatal network correlates with flexible behaviors, but it has remained unclear how activity modulation at these different levels relate to each other during the dynamic transitioning of routing states, leaving open (1) whether neurons show directional interactions in this network, (2) whether directional interactions may relate to network fluctuations and LFP activity states, and (3) whether directional routing states vary with behavioral states.

Here, we addressed these questions by analyzing spiking interactions in the fronto-striatal network in nonhuman primates. We found that the emergence of routing states can be traced back to the sequential temporal lead and lag of single neuron spiking events in relation to multiunit activity (MUA) from distant areas. Overall, spike events in the LPFC preceded (led) MUA increases in the ACC and striatum, while spiking of neurons in the striatum lagged MUA in the LPFC and ACC. The lead/lag relationships defined directional routing states of the LPFC, ACC and striatum network that amplified or switched direction when spikes of single neurons occurred during oscillatory bursts, during attentional shifts, and during decision making.

## Results

We first simulated two reciprocally connected spiking networks to validate a method for quantifying the temporal lead or lag of single neuron spikes relative to multiunit activity at high (<5 ms) temporal precision. In a second step we characterized the lead- and lag- relationships in neuronal ensembles recorded extracellularly from the ACC, LPFC, and striatum of nonhuman primates performing an attention demanding learning task. If not noted otherwise, statistical results on time series are based on randomization tests with extreme-point multiple comparison corrections, and all group-level statistics were corrected for multiple comparisons using false discovery rate (FDR) correction at an alpha level of 0.05.

## Directional interactions between spiking networks

To evaluate how neural activity coordinates between distant networks we simulated two interconnected networks (Izhikevich, 2004) with cell-types and connectivity corresponding closely to LPFC (network 1) and ACC (network 2) (**Figure 1A**, see **STAR Methods** and **Table S2, 3**) (Dombrowski et al., 2001; Medalla et al., 2017). Each network was composed of 1000 spiking neurons and contained 125 recording sites for measuring electrical field dynamics of postsynaptic and transmembrane potentials (see **Figure S1, 2a-c** for an example and network characteristics) (Einevoll et al., 2013). This enabled us to evaluate spike-triggered multiunit activity (stMUA) and the role of LFP oscillatory events on stMUA strength and direction. Within each network, MUAs were near-symmetrically distributed relative to spike events with a zero time-lag (**Figure 1B**). This contrasted to inter-network interactions. Spikes in the simulated LPFC (sLPFC) systematically led MUA in the ACC (mACC, hereafter, the notation sX-mY indicates a pair with spikes *s* from a neuron in area X and multiunit activity *m* in area Y), while sACC lagged mLPFC (**Figure 1B**, permutation test, P<0.05). We quantified the spike lead and lag as the *Lead/Lag index,* which is the difference of stMUA in the 20 ms following versus preceding the spike event. A positive Lead/Lag index value indicates spikes lead MUA, while a negative value shows that spike events lagged MUA. The results from simulations verified that the Lead/Lag index values closely matched the asymmetry of anatomical connectivity strengths between the two networks (**Figure S2D-F**). The Lead/Lag index showed significant non-zero and positive values (spike-lead) for sLPFC-mACC pairs and negative values (spike-lag) for sACC-mLPFC pairs (**Figure 1C**, Wilcoxon test, p<0.001).

**Figure 1.**
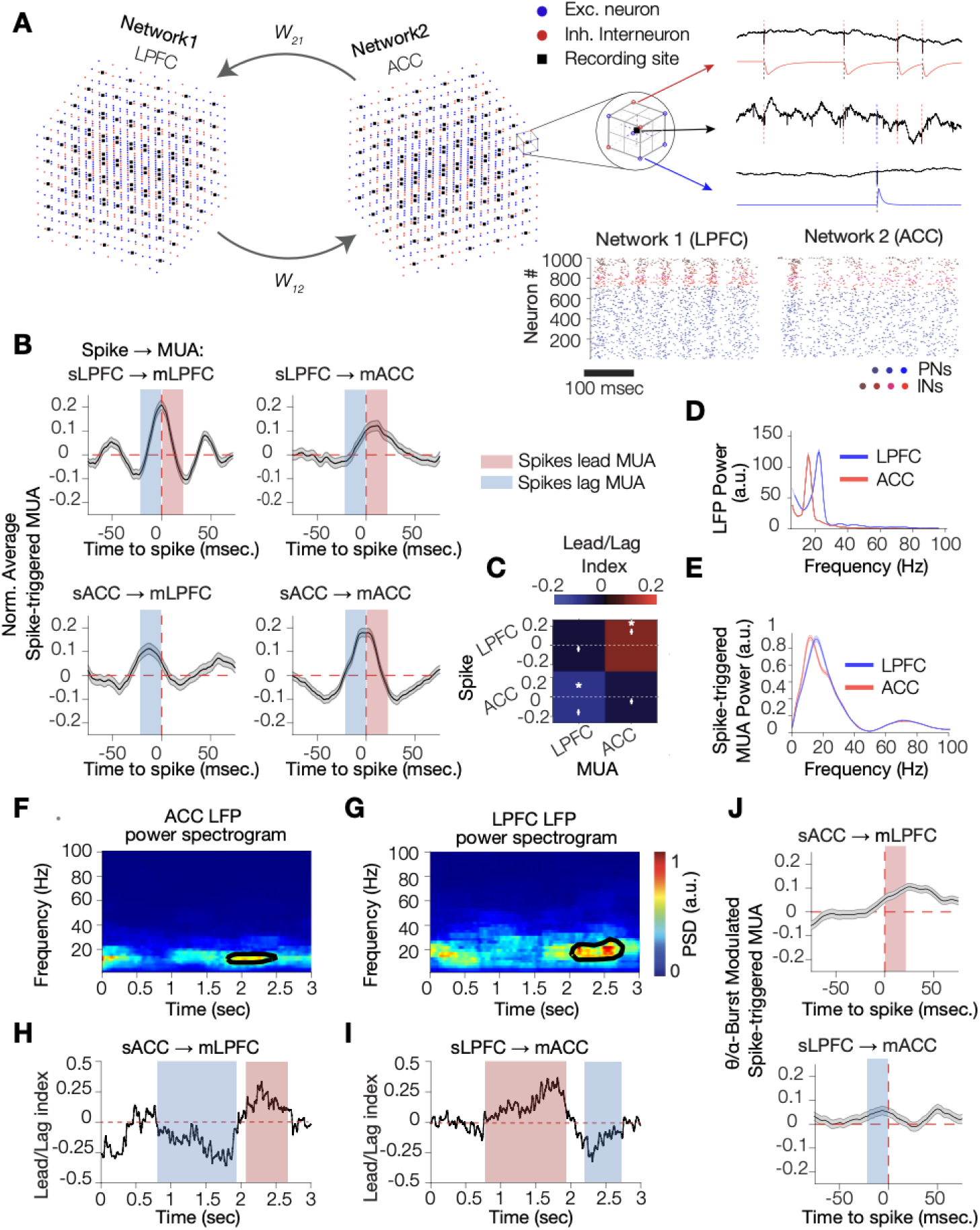
ACC and LPFC spiking network simulations shows systematic inter-network lead and lag temporal orders switching during 8-14 Hz LFP bursts. **(A)** Two networks with excitatory (blue) and inhibitory (red) neurons. Black dots are recording sites in each network measuring the transmembrane and postsynaptic potentials used to extract spiking, multiunit activity (MUA), and local field potentials (LFPs). Raster plots of spiking activity for interneurons (red, INs), and bursting and regular excitatory neurons (blue, PNs). **(B)** Average spike-triggered (st)MUA of the simulated networks 1 (LPFC) and 2 (ACC). Intra-network spikes and MUA show near-symmetric temporal relationships with zero-time lag peaks and oscillatory side-lobes. Inter-network stMUAs have asymmetric shape indicating temporal lead (red shading) and lag (blue shading) relationships. **(C)** Median Lead/Lag inde values across all spike-MUA pairs. LPFC spikes (sLPFC) led MUA in the ACC (mACC) network. White dots an error bars show median ±SE Lead/Lag index value. **(D,E)** Power spectral density (PSD) of LFP **(D)** and of the stMUA **(E)** recorded in each network. **(F)** LFP power spectrogram in the ACC with a theta/alpha burst at ∼2-2.5 sec. Black contours mark LFP burst. **(G)** Example beta burst from LPFC. **(H) Time**-resolved Lead/Lag index values for sACC-mLPFC with a theta/alpha burst occurring at ∼2-2.5 sec. At the time of the burst sACC switched to lea mLPFC. **(I)** Same format as h for mLPFC-sACC. **(J)** The stMUA differences between spikes occurred during versus outside of LFP theta/alpha bursts for sACC-mLPFC (top) and sLPFC-mACC (bottom). Shades indicate significant stMUA in 20 ms preceding (blue) or following (red) spike events.

In each of the simulated networks stMUAs and the LFPs were rhythmically modulated. Spectral power of the LFP and the stMUA showed peaks at ∼20 Hz in the LPFC and at ∼14 Hz in the ACC network (**Figure 1D,E**). The lower frequency in the ACC is due to a higher proportion of inhibitory interneurons with slower synaptic decay compared to the LPFC (Dombrowski et al., 2001; Medalla et al., 2017). LFP oscillations emerged as transient bursts (**Figure 1F,G**). Bursts of increased oscillatory power have been associated with coherent interareal activity and with information routing across brain areas (Palmigiano et al., 2017; van Ede et al., 2018; Zich et al., 2020). Consistent with these suggestions we found that the lead and lag pattern of stMUAs between networks transitioned during oscillatory bursts. Spike events occurring during 14 Hz burst events in the ACC switched from lagging to leading MUA recorded in LPFC (**Figure 1H**), while spikes in LPFC started lagging MUA recorded from ACC (**Figure 1I**). The switch from lagging to leading of spikes in ACC relative to MUA in LPFC during ∼14 Hz bursts was significant (**Figure 1J**, Wilcoxon test, p<0.001), while it was not significant at ∼20 Hz (**Figure S2G**).

## Spikes in ACC and LPFC lead striatum, LPFC spikes lead over ACC

We quantified stMUAs in simultaneous recordings from neurons in the ACC, LPFC, and striatum of nonhuman primates (492, 571 and 534 recording sites, respectively) as detailed before (Oemisch et al., 2019) (**Figure 2A**). We extracted the spike times of isolated neurons and the MUA around each spike time at another electrode recorded in the same area (intra-areal) or a distant area (inter-areal) (**STAR Methods**; **Figure S3** for spike-MUA pair examples). Intra-areal stMUAs showed near-symmetric activity distributions (**Figure 2B**). Compared to the LPFC and striatum, the stMUA in the ACC showed a wider peak and a steeper MUA drop after the spike events (**Figure 2B**, randomization test, P<0.05). Similar to the model predictions (**Figure 1B**), intra-areal stMUAs were rhythmically modulated with the strongest spectral power around 8-14 Hz (theta/alpha) in ACC and 15-25 Hz (beta) in LPFC (**Figure 2C**, randomization test, P<0.05).

**Figure 2.**
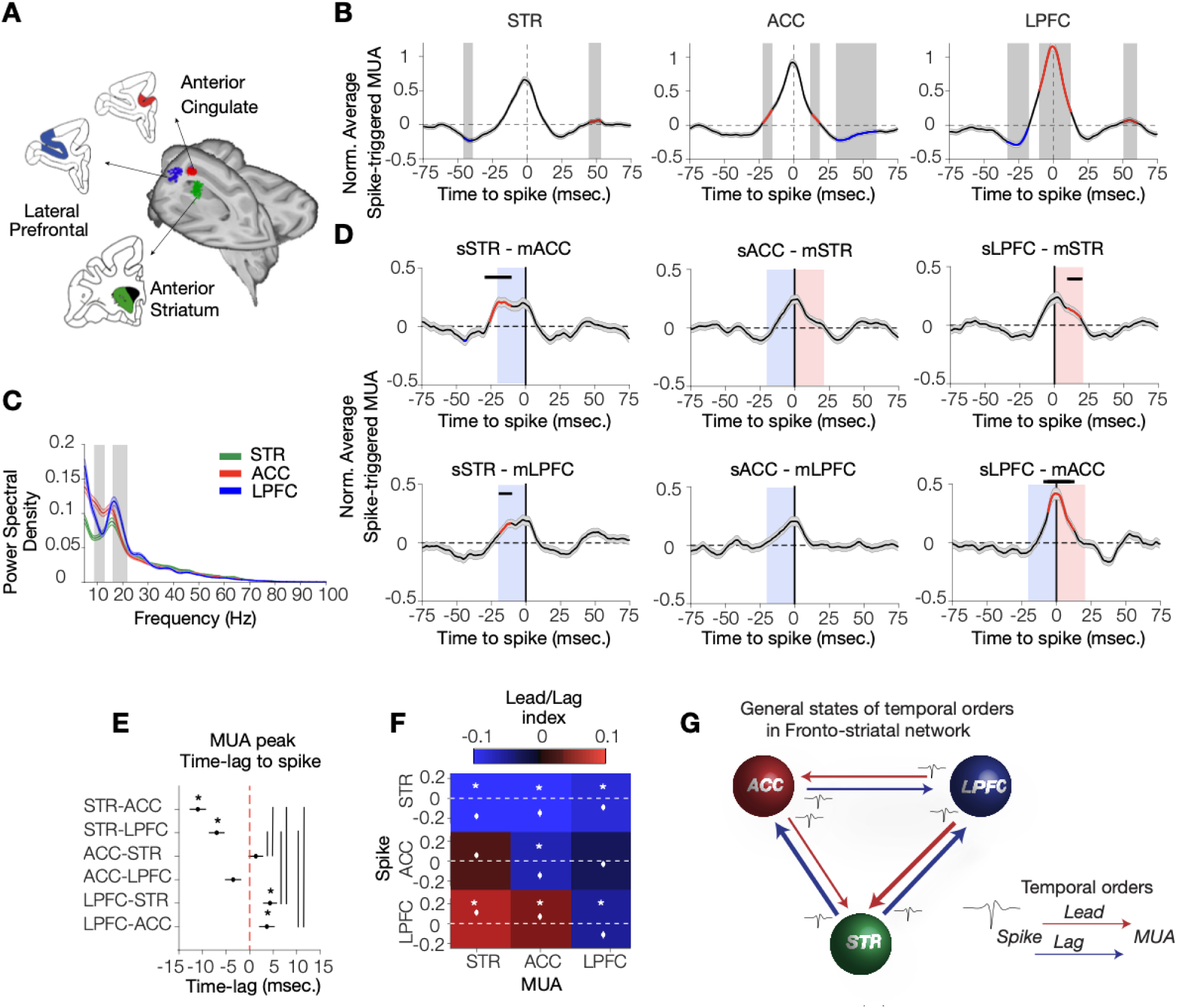
Temporal orders of spiking activity across fronto-striatal areas. (**A)** Neuronal activity was simultaneously recorded from ACC (area 24, red), LPFC (areas 46 and 9, blue), and striatum (caudate and ventral striatum, green). (**B)** Intra-areal stMUA. red/blue signify stMUA values significantly higher/lower than all intra-areal pairs population. (**C)** Power spectral density over stMUA has stronger power peak at ∼16-22 Hz in the LPFC, and at ∼8-14 Hz in the ACC. (**D)** Inter-areal stMUA, Red lines indicate periods when the stMUA of the area pair is significantly higher than the random assignment of areas to the population of spike-MUA pairs (randomization test, P<0.05). Blue/red shades indicate significant non-zero MUA within 20 preceding/following the spike time, respectively. Post-spike MUA was significantly enhanced for spike-MUA pairs: sLPFC-mSTR, sACC-mSTR, sLPFC-mACC at p<0.001; and for sACC-mLPFC at p=0.03, Wilcoxon test, FDR corrected for dependent samples. € Median time lags of the stMUA peaks. Negative/positive values indicate MUA peaked more preceding/followin the spike time (at 0 ms). sLPFC led over mACC and mSTR, mSTR lag behind sACC and sLPFC. Vertical lines show significant pair-wise comparisons (Kruskal-Wallis test, P<0.001, Dunn’s test, P<0.01). Asterisks denote area pairs with time-lags significantly different from zero. (**F)** Lead-Lag index values for all intra-areal (diagonal) and inter-areal (off-diagonal) spike-MUA pairs. sSTR lagged behind mLPFC and mACC, while sLPFC led over mSTR and mACC (Wilcoxon test; p<0.001 for sSTR-mLPFC and sSTR-mACC, p=0.03 for sLPFC-mSTR, FDR correcte for dependent samples). White dots and SE’s show median ± SE. White asterisks denote values significantl different than zero. P-values are corrected for multiple comparisons (see STAR Methods). (**G)** Schematic summar of inter-areal interactions. Red/blue arrows indicate spikes lead/lag MUA, respectively. Line thicknesses reflects lead or lag strengths.

Inter-areal stMUAs had temporal relationships consistent with anatomical data in primates (Ferry et al., 2000; Medalla and Barbas, 2009). On average, sLPFC led mACC and mSTR, sACC tended to lead mSTR, while sSTR lagged both, mACC and mLPFC (**Figure 2D**; randomization test, P<0.05). The red line segments in **Figure 2D** mark the timepoints at which stMUA for the area-pairs was significantly greater than the stMUA from randomized area combinations (randomization test, P<0.05), which included -26 to -9 ms time lag for sACC-mSTR, and -19 to - 8 ms for sLPFC-mSTR (randomization test, P<0.05). The same pattern of results was seen when using as test statistic the time lag with maximal stMUA modulation, which showed that the lead of sLPFC over mSTR and mACC was evident in MUA peaking around +5 and +4.5 ms following the spike event, respectively (**Figure 2E**, Wilcoxon test, P=0.005 and P=0.02 for mSTR and mACC, respectively). In the reverse direction sSTR lagged mACC and mLPFC, with the average stMUA peaks at -12 and -7 ms before the spike, respectively (Wilcoxon test; P<0.001). We confirmed these inter-areal spiking interactions by calculating the Lead/Lag index which also showed a significant lead of sLPFC over mSTR and mACC, as well as a lag of sSTR relative to mACC and mLPFC (**Figure 2F**,**G**; Wilcoxon test; P<0.001, FDR corrected for dependent samples). The lead/lag effects were maximal when using ∼20 ms wide time windows for calculating the Lead/Lag index (**Figure S4**) and were evident in both monkeys (**Figure S5**). The firing rate in all stMUA analyses was controlled by using the same number of spikes for each spike-MUA pair (see **STAR Methods**).

## Spikes in ACC switch during theta/alpha bursts from lagging to leading LPFC

The lead and lag of stMUAs in the model simulations were systematically modulated during oscillatory bursts (**Figure 1H,J**), which raises the question whether the magnitude or direction of empirically measured lead/lag relationships also varied during oscillatory events. We detected reliable bursts in the theta/alpha and beta band in the LFP using a new adaptive width-magnitude thresholding method (**Figure 3A**, see **STAR Methods**, **Figure S6**). Bursts in the theta/alpha and beta bands were most prevalent in the ACC and LPFC, respectively (**Figure S6G,J**). We calculated inter-areal stMUAs separately for spikes occurring during versus outside of LFP bursts. Intriguingly, during theta/alpha bursts, spikes recorded in ACC were followed by stronger sACC-mLPFC stMUA amplitudes than spikes outside of the burst and switched from a net lag (**Figure 2F**) to a net lead over MUA in mLPFC (**Figure 3B**, randomization test, P<0.05). Spikes in ACC also significantly increased the amplitude of its lead over MUA in STR during theta/alpha bursts (**Figure 3B**, randomization test, P<0.05). Consistent with this finding, spikes during theta/alpha bursts in the striatum (sSTR) showed an amplified lag relative to MUA from LPFC and ACC (**Figure 3B**, randomization test, P<0.05). In the LPFC spike events during versus outside of theta/alpha bursts were preceded and followed by stronger MUA in the ACC and striatum (**Figure 3B**, randomization test, P<0.05, red line segments indicate the timepoints at which in burst stMUA for the area-pairs was significantly greater than the in burst stMUA from randomized area combinations; randomized permutation test, P<0.05). Calculating the Lead/Lag indices across area-pairs confirmed that during theta/alpha bursts the lead was amplified for sACC-mLPFC pairs, and the lag was amplified for sLPFC-mACC pairs (**Figure 3C**, Wilcoxon test, P=0.007 and P=0.036, respectively).

**Figure 3.**
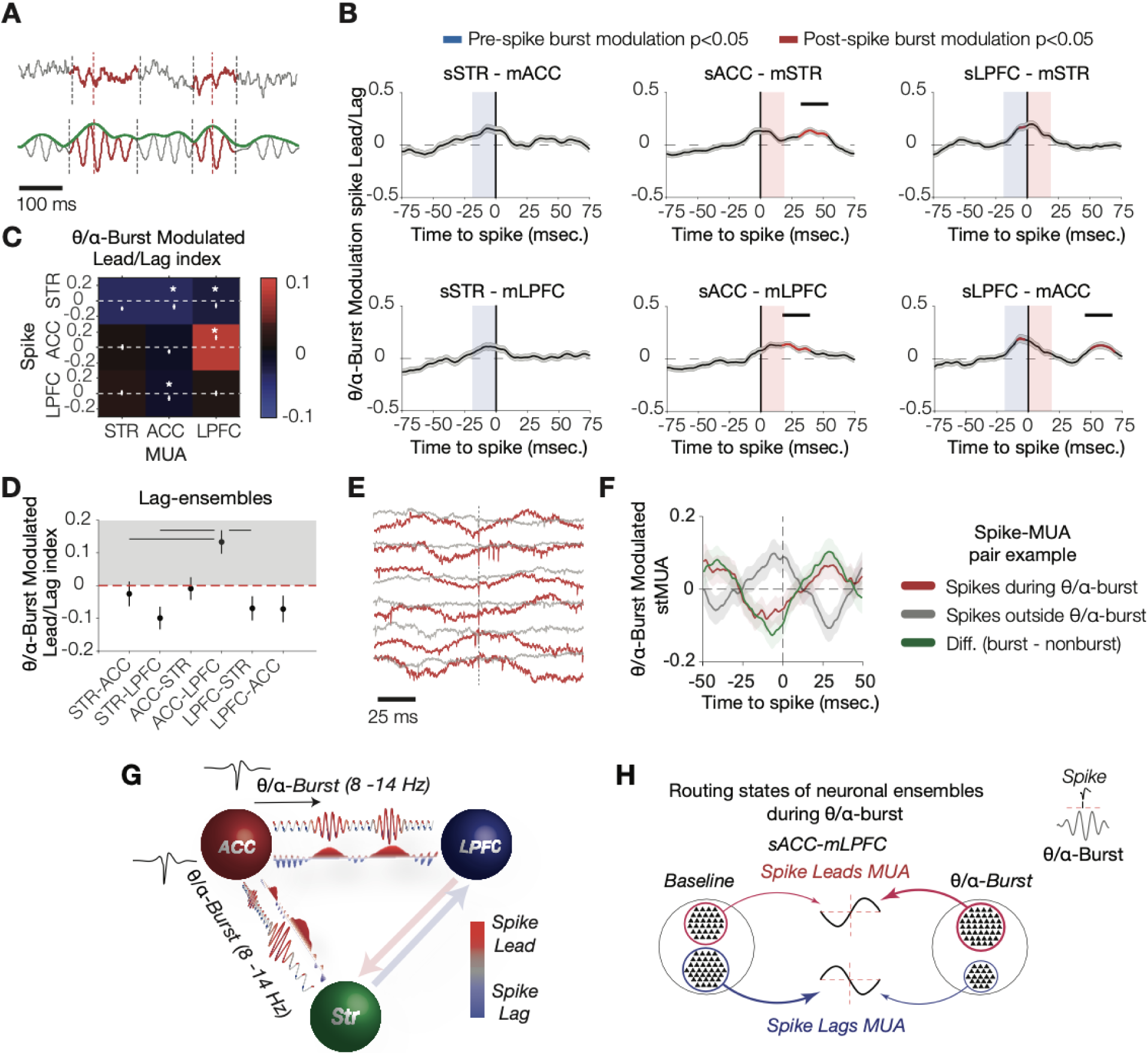
ACC spikes switch to lead over LPFC multiunit activity during Theta/alpha (8-14 Hz) bursts. **(A)** A wideband LFP example segment with 8-14 Hz bursts in red (top) and bandpass filtered signal (bottom) wit envelope in green. **(B)** The stMUA difference for spikes inside versus outside of theta/alpha bursts for area-pair combinations. Red lines indicate periods where in burst stMUA is significantly higher the area-pair than the randomized population. Red and blue shades indicate significant non-zero stMUA difference of theta/alpha burst versus non-burst within 20 ms following/preceding spikes, respectively. **(C)** Similar Lead-Lag index adjacenc matrix as in (**Figure 2f**) for stMUA differences of spikes during versus outside of theta/alpha bursts. White dots an SE’s show median ± SE. White asterisks denote values significantly different than zero. **(D)** Lead/Lag index values flipped signs during theta/alpha bursts in the sACC-mLPFC lag-ensembles (Kruskal Wallis test, P=0.01, Wilcoxo test, P=0.03 for nonzero ACC-LPFC. Black horizontal lines show significant differences between area-pairs). **(E)** LFP segments for a spike-MUA pair example around spikes (±75 ms), for spikes inside (red) versus outside (grey) of the theta/alpha bursts. **(F)** The spike-MUA pair example in (**E**) switched from a lag to a lead state. Red, grey, an green colors denote stMUAs inside bursts, outside bursts, and their difference, respectively. **(G)** Schematic summary of theta/alpha burst modulated inter-areal interactions. The filled red/blue portion of the lines indicates where spikes lead and lag relative to theta/alpha bursts. **(H)** Schematic summary of lag to lead switch in the sACC-mLPFC during theta/alpha burst.

We next asked whether the burst-dependent switch and amplification of directional interactions were based on stMUA modulation in leading or lagging spike-MUA pairs. Ensembles of leading and lagging spike-MUA pairs were extracted by projecting the stMUA profiles of individual interareal spike-MUA pairs to a principal components (PCs) space (**Figure S7A,B**, see **STAR Methods**). Across PCs, only one PC (PC2) showed an asymmetric stMUA temporal profile (**Figure S7C,D**). The asymmetric PC profile suggested that PC2 captured the directional component of inter-areal interactions, which we corroborated by finding that stMUAs with positive and negative PC2 loading had leading and lagging stMUA profiles, respectively (**Figure S7E**). This finding enabled using the sign of the PC2 loading to group spike-MUA pairs as members of a *lead-ensemble* when they had a negative loading on the PC2, and conversely to group spike-MUA pairs as members of a *lag-ensemble* when they showed a positive PC2 loading (See **Table S1** for details on the number of spike-leading and spike-lagging spike-MUA pairs for each brain area). We found that the switch from lagging to leading of sACC over mLPFC was largely due to spike-MUA pairs of the lag-ensemble switching to the lead-ensemble during theta/alpha bursts (**Figure 3D**, Kruskal Wallis test, P=0.002, Wilcoxon test for non-zero modulation P<0.001). The stMUA directionality of lead-ensembles was unchanged during theta/alpha bursts across area-pairs (**Figure S7F,G**). **Figures 3E** and **F** illustrates an example sACC-mLPFC pair with stMUA that switched from lagging outside of bursts (grey) to leading during theta/alpha bursts (red). In summary, during theta/alpha bursts, spikes in ACC transitioned in a routing state with LPFC and STR from lagging to leading relative to LPFC and to a stronger lead over STR (**Figure 3G**). The lag-to-lead switch was largely based on spike- MUA pairs of the lag-ensemble joining the lead-ensemble during theta/alpha bursts (**Figure 3H**).

## Lead of LPFC over striatum is amplified during beta bursts

Similar to theta/alpha, we detected (15-25 Hz) beta bursts in the LFP of each area and quantified inter-areal stMUAs for spikes occurring during beta bursts (**Figure 4A**, **Figure S6D**,**H-J**). Comparing stMUAs during beta bursts to non-burst periods showed that during beta bursts, the lead of sLPFC over mSTR was amplified and, reciprocally, sSTR more strongly lagged mLPFC and mACC (**Figure 4B**, Wilcoxon test, P<0.001). Consistent with this finding we found a significant difference of the Lead/Lag index for burst versus non-burst periods for stMUA of sLPFC-mSTR pairs (**Figure 4C**, p<0.001, Wilcoxon test). Similar to the theta/alpha burst results, the amplified lead of LPFC was largely due to spike-MUA pairs that lagged outside of bursts but switched to lead during beta bursts (**Figure 4D**, **Figure S7H**,**I**). This switch from a lag- to a lead- state during beta bursts was evident at the level of individual spike-MUA pairs as shown for an example in **Figure 4E**,**F** (for more examples see **Figure S8**). In summary, the neuronal spike output of LPFC amplified its net lead over MUA in the striatum during beta bursts (**Figure 4G**). This effect that was driven by neurons of the lag-ensemble joining the lead-ensemble during beta bursts (**Figure 4H**). The same pattern of results was evident in both monkeys (**Figure S9**).

**Figure 4.**
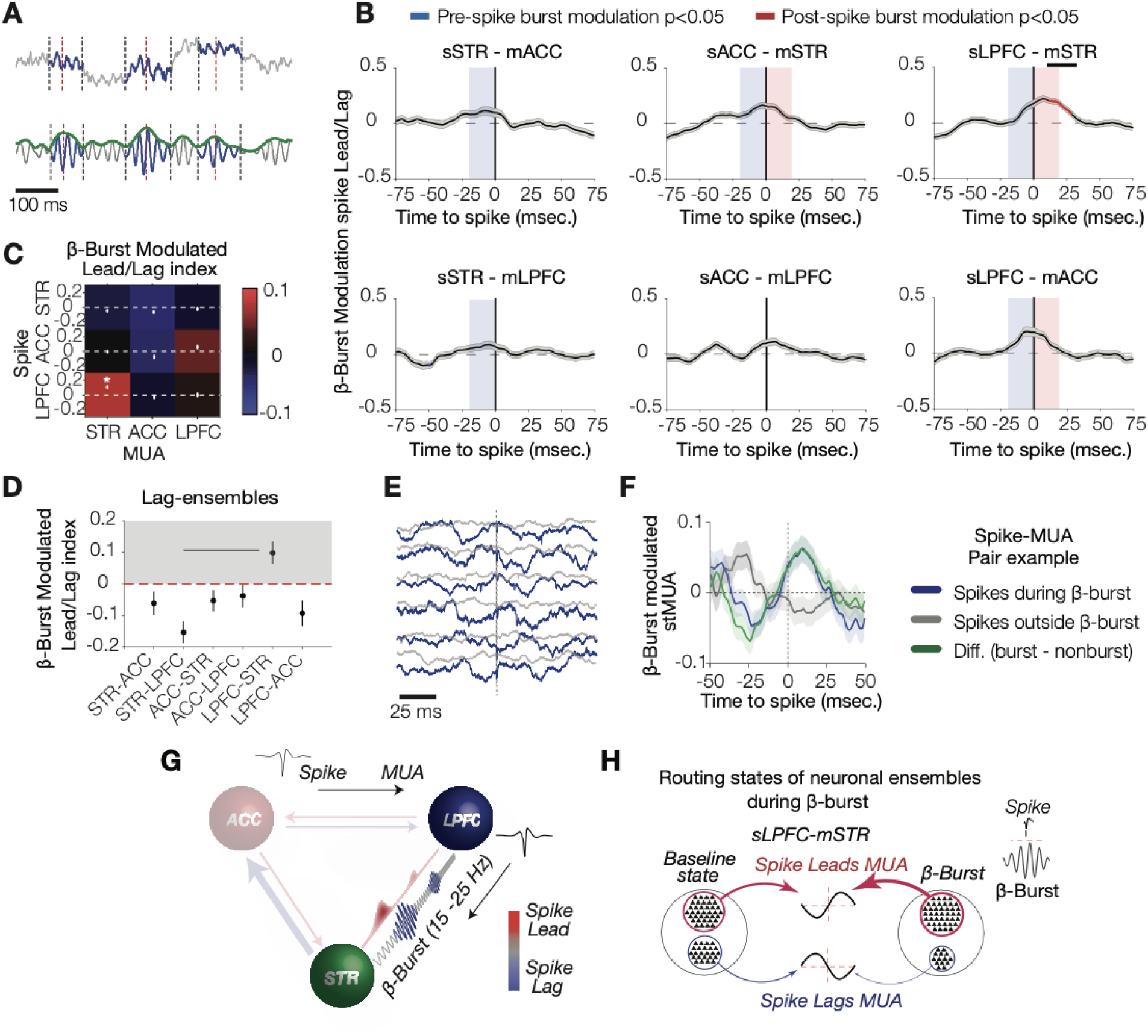
LPFC spikes amplified their lead over striatum during beta (15-25 Hz) bursts. **(A)** A wideband LFP example segment with beta bursts in blue (top) and bandpass filtered signal (bottom) with envelope in green. **(B)** Average differences of stMUA’s inside versus outside of beta bursts for area-pair combinations. Red lines indicate periods where in burst stMUA is significantly higher the area-pair than the randomized population. Red and blue shades indicate significant non-zero stMUA difference of beta burst versus non-burst within 20 ms following/preceding spikes, respectively. **(C)** Similar Lead-Lag index adjacency matrix as in (**Figure 3C**) for stMUA differences of spikes during versus outside of beta bursts. White dots and SE’s show median ± SE. Whit asterisks denote values significantly different than zero. **(D)** Lead/Lag index values was flipped for lag-ensembles in the sLPFC-mSTR during beta bursts. Black horizontal lines show significant differences between area-pairs. **(E)** LFP segments for a spike-MUA pair example around spikes (±75 ms), for spikes inside (blue) versus outside (grey) of the beta bursts. **(F)** Beta burst modulated stMUA for the spike-MUA pair example in (**E**) for spikes inside (blue) and outside (grey) of the bursts, and their difference (green). **(G)** Schematic summary of beta burst modulated inter- areal interactions. The filled red/blue portion of the lines indicates where spikes lead and lag relative to beta bursts. **(H)** Schematic summary of lead amplification in the sLPFC-mSTR during beta burst.

## Theta/alpha-tuned lead-ensemble in ACC responds to attention cue

We next asked whether the directional spiking interactions of neuronal ensembles changed during selective attention states. Neural activity was recorded while monkeys used the color onset of peripheral stimuli as attention cue to covertly shift attention left or right from a central fixation point, and subsequently used the motion direction of the attended peripheral stimulus to plan a saccadic choice up or down from the fixation point (**Figure 5A**) (Oemisch et al., 2019). The saccadic choice was triggered a few hundred milliseconds later by a transient dimming of the attended stimulus. We analyzed stMUA’s for spike events occurring in a 0.5 s window following the attention cue onset and found that spikes of ACC neurons showed significantly amplified lead over MUA in the striatum during this attention period (**Figure 5B**, **Figure S10A**). This attention modulation of the lead was evident at 12-30 ms time lag for sACC-mSTR pairs of the lead-ensemble, at 4-27 ms time lag for the sACC-mSTR pairs of the theta/alpha-tuned lead- ensemble, and at 15-25 ms time lag when considering all sACC-mSTR spike-MUA pairs (randomization tests, all P<0.05; **Figure 5B**; **Figure S10A**). Notably, however, for the ACC connection to LPFC (sACC-mLPFC pairs), the amplified lead during the attention period was limited to the stMUA pairs of the theta/alpha-lead-ensembles with a significant stMUA difference at 7-22 ms time lag (randomization test, P<0.05; **Figure 5B**). We quantified the attention effect as the difference of the Lead/Lag index values during the attention cue period versus irrespective of that period, which confirmed that on average neuronal spiking in the ACC amplified its lead over the striatum following the onset of the attention cue (Wilcoxon test, P=0.05; **Figure 5C**). Attention also amplified the lead for the subset of sACC-mSTR pairs that showed an ACC lead over the striatum (Wilcoxon test, P=0.03; **Figure 5D**) and for sACC-mSTR pairs that led the striatum during theta/alpha-bursts (Wilcoxon test, P=0.03; **Figure 5E**). Analyzing the Lead/Lag index also confirmed that attention exclusively amplified the lead of the theta/alpha-tuned lead-ensemble of sACC over mLPFC (Wilcoxon test, P=0.03; **Figure 5E**; **Figure S10**).

**Figure 5.**
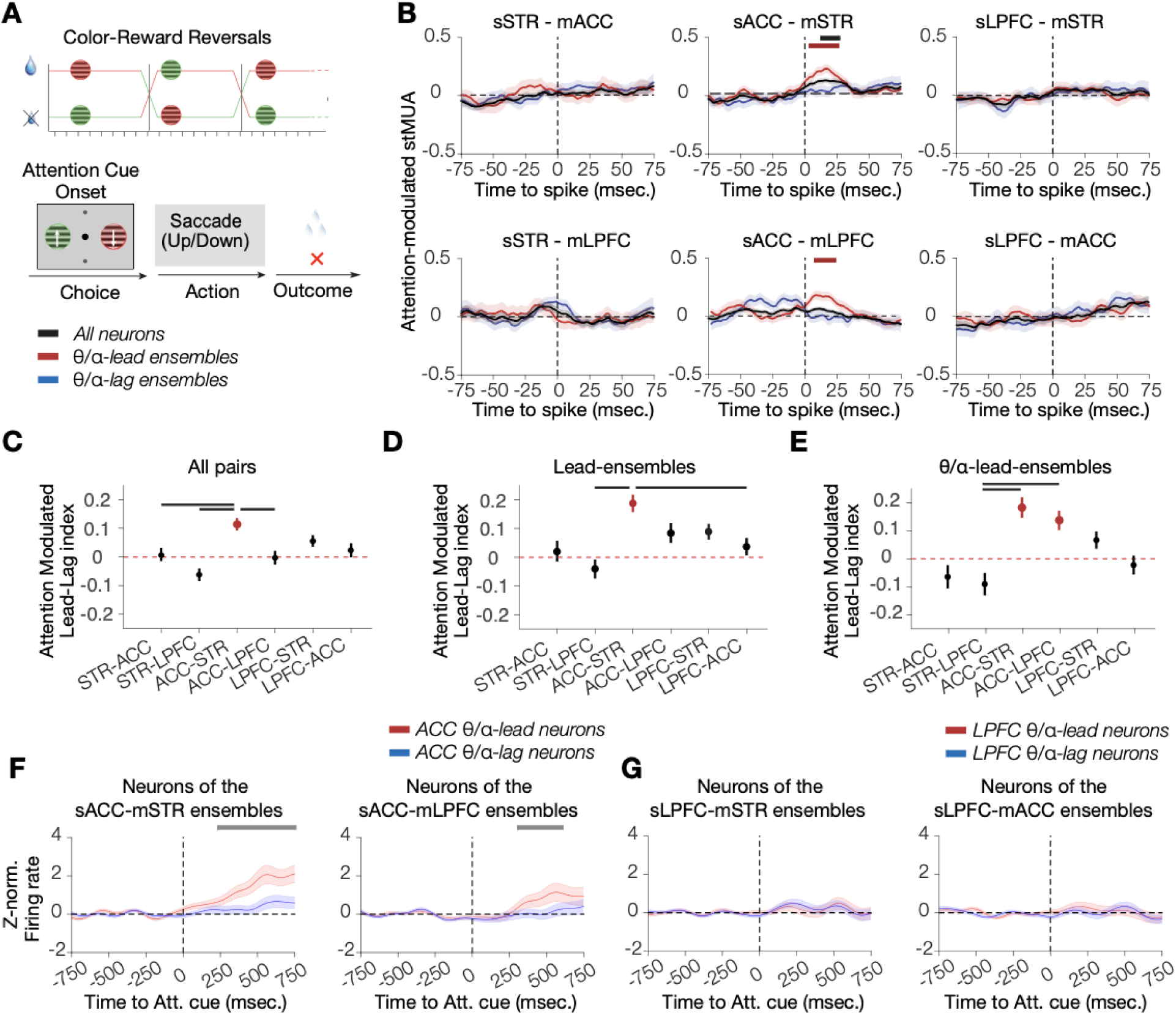
Theta/alpha-subnetworks of neurons in the ACC route attentional states. **(A)** Monkeys performed a task requiring covertly attending a peripheral stimulus and making a saccadic response to a transient luminance change of the attended stimulus to receive reward (see **STAR Methods**). The stimulus color-reward associatio reversed after ∼30 trials**. (B)** Attention-modulated stMUA for all neurons (black), theta/alpha-lead (red) an theta/alpha-lag (blue) neuronal ensembles. The sACC showed stronger lead over mSTR in response to an attentio cue on a population level and over mLPFC only in theta/alpha-lead-ensembles. Black/red horizontal lines show significant nonzero attention-modulated stMUA for all neurons and theta/alpha-lead neuronal ensembles, respectively. (**C-E)** Attention-modulated lead/lag index for all neurons **(C)**, lead-ensembles **(D)**, and theta/alpha- lead-ensembles **(E).** Bars in red are significantly different than zero (Kruskial Wallis test, P<0.01 for all three, an Wilcoxon test, P<0.05, FDR corrected, see **STAR Methods**). Black horizontal lines denote pairs with significantly different lead/lag index**. f,** Neuronal spike rate dynamics in response to attention cue for (**F**) the ACC neurons identified by their temporal orders during theta/alpha bursts relative to the mSTR (left) and the mLPFC (right). **(G)** Same as in (**F**) but for LPFC neurons identified by their temporal orders during theta/alpha bursts relative to the mSTR (left) and the mACC (right). The theta/alpha-lead and theta/alpha-lag subnetworks are shown in red and blue respectively. gray horizontal lines denote time periods were the two subnetworks show different rate response dynamics (permutation test, P<0.05, see **STAR Methods**). Overall, the ACC neurons identified by their theta/alpha- lead over striatum showed earlier onset response compared with the LPFC. Error bars denote standard error of the mean.

We next asked whether neurons of stMUA pairs of the lead and lag-ensembles show differential firing rate dynamics after the attention cue onset. We next computed the peri-stimulus time histogram (PSTH) around the time of the attention cue for neurons labeled either as lead or lag based on their stMUA profiles during theta/alpha bursts. In the ACC, neurons identified by a theta/alpha-specific lead showed larger and distinct firing rate dynamics than ACC neurons of stMUA pairs that lagged during theta/alpha bursts (**Figure 5F**.) The firing rates of neurons with a lead versus lag relationship during theta/alpha bursts were significantly different for sACC- mSTR pairs at 240-750 ms and for sACC-mLPFC pairs at 360-610 ms following attention-cue onset (permutation tests, P<0.05). The group of neurons in ACC that showed a theta/alpha-band specific lead relation can be considered to constitute an *attention modulated theta/alpha-lead subnetwork*. In contrast to the attentional effect of this theta/alpha-lead subnetwork, neurons identified by their lead or lag during beta bursts did not show different attentional effects on their firing rate dynamics (**Figure S10C,D**).

## ACC spiking lead LPFC and striatum during a choice

So far, we found that ACC neurons of the theta/alpha-lead subnetwork amplified their lead over LPFC and striatum in response to an attention cue. A different change of routing states occurred during the preparation of the choice immediately prior to the saccadic eye movement. In response to the transient dimming of the peripherally attended stimulus, the stMUAs with neurons from both, LPFC and ACC, showed an amplified lead over MUA in the striatum (**Figure 6A**). This amplified lead was evident when considering all spike-MUA pairs (the overall population level), when considering the subgroup of spike-MUA pairs that led the striatum (the level of a leading directional subnetwork), and - with a later onset – when considering pairs of ACC or LPFC neurons that lagged MUA in the striatum (time periods with significantly amplified lead are shown in **Figure 6A**; for sACC-mSTR pairs: at 9-19 ms and 46-60 ms time lag in the lead-ensembles and at 54-70 ms time lag in the lag-ensembles; for sLPFC-mSTR pairs: at 12-75+ ms time lag in the lead-ensembles, and at 25-51 ms time lag in the lag-ensembles, randomization test, P<0.05). In addition, during the choice period, the lead of stMUAs with spikes from ACC and MUA in LPFC was amplified in the lead-ensembles (at 22-41ms time lag, randomization test, P<0.05; **Figure6A**). We confirmed the finding of an amplified lead during the choice by computing the Lead/Lag index values for all sACC-mSTR and sLPFC-mSTR stMUA pairs (Wilcoxon test, P=0.03 and P=0.01, respectively; **Figure 6B,C**), and for sACC- mSTR and sLPFC-mSTR stMUAs pairs of the lead-ensembles (Wilcoxon test, P<0.01, for both area pairs, **Figure 6C**). In addition, the Lead/Lag index showed a significantly amplified lead of sSTR-mLPFC stMUAs of the lead-ensemble (Wilcoxon test, P=0.016, **Figure 6C**).

**Figure 6.**
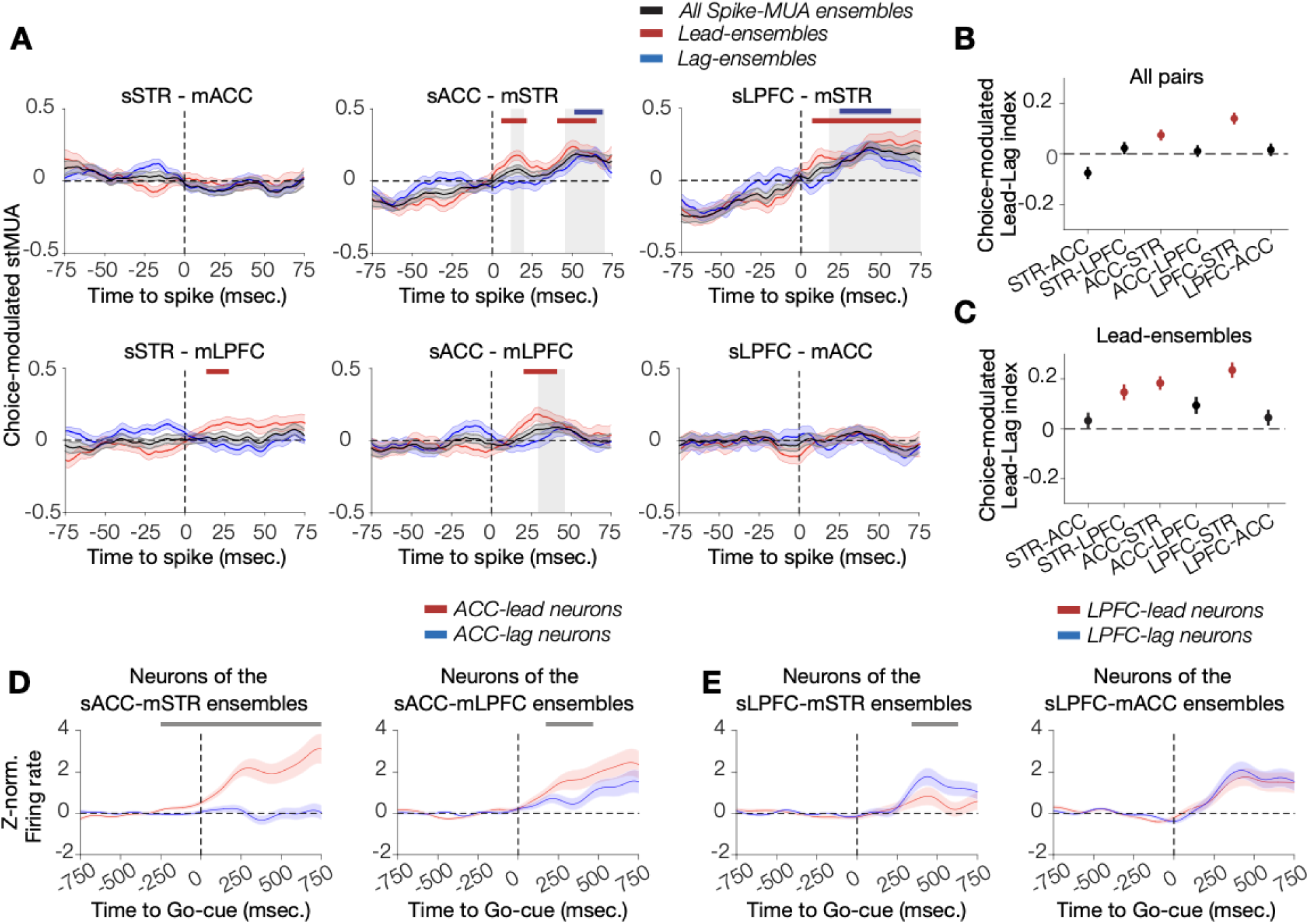
The ACC predominates the fronto-striatal network response to a choice. **(A)** Choice-modulated stMUA for all neurons (black), lead (red) and lag (blue) neuronal ensembles. The sACC and sLPFC show stronger lead over mSTR during a choice on a population level and both the lead- and lag-neuronal ensembles. The sACC showed stronger lead over mLPFC on a population level and the lead-neuronal ensembles (randomizatio test, P<0.05, see **STAR Methods**). Horizontal lines denote significant nonzero stMUA for all neurons (black), lead (red) and lag (blue) neuronal ensembles, indicating earlier lead-onset for lead neuronal ensembles. **(B,C)** Choice-modulated lead/lag index for all neurons **(B)**, and lead-ensembles **(C).** Bars in red are significantly different than zero. Black horizontal lines denote pairs of significantly different lead/lag index (Kruskial Wallis test, P<0.01 for all two, and Wilcoxon test, P<0.05, FDR corrected, see **STAR Methods**)**. (D),** Neuronal spike rate dynamics in response to Go-cue for the ACC neurons identified by their temporal orders relative to the mSTR (left) and the mLPFC (right). **(E)** Same as in (**D**) but for LPFC neurons identified by their temporal orders relative to the mSTR (left) and the mACC (right). The lead/lag subnetworks are shown in red/blue, respectively. Gray horizontal lines denote time periods were the two subnetworks showed different rate response dynamics (permutation test, P<0.05, see **STAR Methods**). Overall, the ACC neurons identified by their lead over striatum showed earlier onset response compared with the LPFC. The LPFC showed stronger response dynamics in the neurons that lagged the striatum. Error bars denote standard error of the mean.

Similar to the analysis of attention, we asked whether neurons belonging to lead- and lag- ensembles responded differently during a choice. We found that the ACC neurons that led over the MUA in the striatum and LPFC showed on average stronger and distinct firing rate dynamics than neurons with spikes lagging MUA (**Figure6D**). The firing response difference was significant at -270-750+ ms and 210-430 ms after the onset of the choice-triggering dimming of the peripheral stimulus for neurons identified by their lead in the sACC-mSTR and sACC- mLPFC ensembles, respectively (permutation tests, both P<0.05). In LPFC, neurons that lagged MUA in the striatum showed distinct and stronger firing responses during the choice than neurons leading MUA in the striatum (**Figure 6E**, 350-680 ms after the dimming choice event, permutation test, P<0.05). Neurons of lead and lag-ensembles of sLPFC-mACC pairs that their lead or lag were not modulated during functional states did not show different firing rate modulation during the choice (**Figure6E**).

## Discussion

Here, we quantified the spatio-temporal dynamics of interareal spiking interactions and found that directional routing states emerged within ∼20 ms time windows. Despite sparse anatomical connectivity of medial and lateral prefrontal cortex to the striatum (Ferry et al., 2000; Morris et al., 2016), our methodology identified lead and lag activity patterns between these brain areas contiguous with their known anatomical connectivity (**Figure 2**)(Ferry et al., 2000; Haber and Knutson, 2010). On average, spikes in the LPFC led MUA in the ACC and striatum, while spikes in the striatum lagged MUA in the ACC and LPFC (**Figure 2**). The fast ∼20 ms timescale characterization of these directional spiking interactions in the fronto-striatal network is new for both, rodents and non-human primates. In rodents cross-areal, fronto-striatal cell assemblies with fast temporal coordination have been recently described, but without elucidating the directionality of spiking interactions (Oberto et al., 2021). In primates, previous studies succeeded to quantify neuronal interactions but with time resolutions of hundreds of milliseconds and across populations of neurons (Antzoulatos and Miller, 2011; Oemisch et al., 2015). Thus far, directional networks have been reported mostly in sensory brain areas using spiking activity patterns of groups of neurons (Semedo et al., 2019; van Kempen et al., 2021), or in higher-order brain areas by inferring directional neuronal communication using LFP measures (Gregoriou et al., 2009; Salazar et al., 2012; Antzoulatos and Miller, 2014; Vezoli et al., 2021). The results of our study not only quantified lead and lag coordination of spiking activity at a fast <20 ms timescale across distant higher order brain areas, but also provide direct insight about the contributions of single neurons in forming these directional routing states.

We found that fast timescale spiking interactions were routed through frequency specific channels that were evident in the LFP dynamics (**Figure 1F-J**). The LPFC led striatal activity stronger when its spike occurred during 15-25 Hz beta bursts (**Figure 4A-C**). In the ACC, spikes that occurred during 8-14 Hz theta/alpha bursts switched their directional relation to lead MUA in the LPFC (**Figure 3A-C**). Both, the amplification and switching effects were primarily driven by spike-lagging neuronal ensembles switching direction during LFP bursts (**Figure 3D**,**4D**). This result indicates that the transitioning between routing states is likely determined by modulating the timing of outgoing neural activities in spike-lagging neurons within their local population into a more synchronized state with spike-leading neurons. The network simulation showed that spectral peaks of LFP activity matched spectral peaks of the stMUA profiles, suggesting that oscillatory bursts more likely occur at frequencies that correspond to the resonance frequency of outgoing neuronal spiking activities (**Figure 1D**,**E**). These resonance frequencies of outgoing spiking activities reflect synaptic time decays in each senders’ brain area (Hahn et al., 2014). During oscillatory bursts, spikes occur more synchronized at the resonance frequency of their local population which as a consequence enhances the efficacy to activate postsynaptic neurons in a recipient area (Kumar et al., 2008; Hahn et al., 2014), and facilitates successful information routing between brain areas through frequency specific channels, as previously shown in models (Palmigiano et al., 2017; Hahn et al., 2019) and at the level of LFP-LFP interactions(Canolty et al., 2010).

Our study suggests that the ACC plays a significant role in mediating the routing of information during functionally important events. We found that the directionality of spiking interactions changed transiently during selective attention and decision-making. This functional specificity of the routing states might reflect a hierarchical organization of fronto-striatal network communication. The ACC predominated the network by amplifying its lead over the LPFC and striatum during both decision-making and selective attention (**Figure 5E,6C**). In contrast, the lead of LPFC spiking output was only amplified during decision making and over the striatum (**Figure 6A-C**). These results illustrate a functional hierarchy at the level of single neuron communication states. In this hierarchy, the ACC ranks higher than the LPFC, and both rank higher than the striatum consistent with previous studies of hierarchical network organization (Haber and Knutson, 2010; Wang, 2022). According to this view, fronto-striatal network interactions do not proceed with a default lead or lag relation, but different nodes of the network can dynamically gain modulate the temporal lead of the network depending on the functional states. The transient amplification in the lead of ACC output reflects its higher rank in the network hierarchy and is consistent with a role of the ACC in top-down control of the PFC when attention is allocated, prediction errors are evaluated, or uncertainty is estimated to determine the degree of top-down control (Shenhav et al., 2013; Womelsdorf and Everling, 2015; Banaie Boroujeni et al., 2021). The functionally transient, rather than persistent ‘control state’ may be related to the stronger inhibitory activity of the ACC (both intra- and interareal inhibitory control)(Medalla and Barbas, 2009; Medalla et al., 2017), which imposes high biological costs for this area to exert its function. Such transient control states – as opposed to stable, or ’persistent’ control states - might facilitate the flexible regulation of routing states depending on rapidly changing task demands (Shenhav et al., 2013; Aben et al., 2020; Boroujeni et al., 2022).

Lastly, we showed that the functional modulation of the lead and lag relation of a neuron’s spike within ∼20 ms time windows predicted the firing rate response at a slower ∼150 ms timescale. We found that neurons with functionally gain modulated spike-lead over distant MUA (i.e. the theta/alpha-leading ensembles during selective attention, and lead-ensembles during a choice) showed stronger and distinct firing rate dynamics of functional responses when compared with neurons from lag-ensembles (**Figure 5F,6D**). These results imply that variable timescales of information processing previously associated with different brain areas (Murray et al., 2014; Luczak et al., 2015; Wang, 2022) can be more precisely understood in a relative framework that quantifies the temporal relation of spiking activity in one brain area relative to another brain area. In such a relative framework, rather than analyzing aggregated functional responses of groups of neurons in individual brain regions, the precise, cross-areal temporal sequencing of neuronal activity is the central unit of functional relevance.

Together, our results demonstrate, to our knowledge for the first time the spatio-temporal dynamics of spiking activity interactions across multiple spatial and temporal scales and between distant brain areas during complex cognitive functions. We have shown that fast directional spiking interactions across distant brain areas instantiate routing states that dynamically switch or amplify during coherent local population activity and when the fronto-striatal network contributes to selective attention and decision-making. Such fast spatiotemporally characterized dynamics predict neuronal activity at slower timescales during relevant functional states.

## Author Contributions

K.B.B. and T.W. conceived the research. K.B.B. developed the methodology. K.B.B. conceptualized and performed modeling, analysis, and visualization. T.W. Supervision. K.B.B. and T.W. wrote the paper.

## Data and materials availability

Code and analysis pipelines will be publicly available upon publication.

## Figure Legends

## Acknowledgments

This work was supported by the National Institute of Mental Health (R01MH123687), by the National Institute of Biomedical Imaging and Bioengineering (R01EB028161), and by the Canadian Institutes of Health Research CIHR (Grant MOP_102482). The funders had no role in study design, data collection and analysis, the decision to publish, or the preparation of this manuscript.

